# Protein QTL analysis of IGF-I and its binding proteins provides insights into growth biology

**DOI:** 10.1101/2020.03.25.007724

**Authors:** Eric Bartell, Masanobu Fujimoto, Jane C. Khoury, Philip R. Khoury, Sailaja Vedantam, Joel Hirschhorn, Andrew Dauber

**Affiliations:** Department of Genetics, Harvard Medical School, Boston, MA 02115, USA; Broad Institute of MIT and Harvard, Cambridge, MA, USA; Cincinnati Center for Growth Disorders, Division of Endocrinology, Cincinnati Children’s Hospital Medical Center, University of Cincinnati College of Medicine, Cincinnati, OH 45229, USA; Division of Pediatrics and Perinatology, Tottori University Faculty of Medicine, Yonago, Tottori 683-8504, Japan; Division of Biostatistics and Epidemiology, Cincinnati Children’s Hospital Medical Center, University of Cincinnati College of Medicine, Cincinnati, OH 45229, USA; Heart Institute Research Core, Cincinnati Children’s Hospital Medical Center, University of Cincinnati College of Medicine, Cincinnati, OH 45229, USA; Division of Endocrinology, Boston Children’s Hospital, Harvard Medical School, Boston, MA 02115, USA; Division of Endocrinology, Children’s National Hospital, Washington, DC 20010, USA; Department of Pediatrics, George Washington University School of Medicine and Health Sciences, Washington, DC 20052

## Abstract

The growth hormone and insulin-like growth factor (IGF) system is integral to human growth. Genome-wide association studies (GWAS) have identified variants associated with height and located near the genes in this pathway. However, mechanisms underlying these genetic associations are not understood. To investigate the regulation of the genes in this pathway and mechanisms by which regulation could affect growth, we performed GWAS of measured serum protein levels of IGF-I, IGFBP-3, PAPP-A2, IGF-II, and IGFBP-5 in 839 children (3-18 years) from the Cincinnati Genomic Control Cohort. We identified variants associated with protein levels near *IGFBP3* and *IGFBP5* genes, which contain multiple signals of association with height and other skeletal growth phenotypes. Surprisingly, variants that associate with protein levels at these two loci do not colocalize with height associations, confirmed through conditional analysis. Rather, the *IGFBP3* signal (associated with total IGFBP-3 and IGF-II levels) colocalizes with an association with sitting height ratio (SHR); the *IGFBP5* signal (associated with IGFBP-5 levels) colocalizes with birth weight. Indeed, height-associated SNPs near genes encoding other proteins in this pathway are not associated with serum levels, possibly excluding PAPP-A2. Mendelian randomization supports a stronger relationship of measured serum levels with SHR (for IGFBP-3) and birth weight (for IGFBP-5) than with height. In conclusion, we begin to characterize the genetic regulation of serum levels of IGF-related proteins in childhood. Furthermore, our data strongly suggest the existence of growth-regulating mechanisms acting through IGF-related genes in ways that are not reflected in measured serum levels of the corresponding proteins.

## Introduction

The IGF system is known to play a role in growth (Ranke 2018 1) and disruption of the genes encoding growth hormone (*GH1*), insulin-like growth factor-1 (*IGF1*), or insulin-like growth factor-2 (*IGF2*) which leads to a variety of growth disorders (Storr 2019 2, David 2011 3, Savage 2011 4). In mice, knockout of *Igf1, Igf2*, or the gene encoding the IGF-I receptor (*Igf1r*) reduces birth weight and body size: birth weight is reduced 40% and 60% for *Igf1* and *Igf2* knockouts, respectively (Liu 1993 5, Powell-Braxton 1993 6, DeChiara 1990 7, Baker 1993 8, Sferruzzi-Perri 2011 9). Furthermore, overexpression of *Igf2* results in overgrowth (Baker 1993 8, Sferruzzi-Perri 2011 9). In humans, total IGF-I and total IGFBP-3 are used as clinical biomarkers in the assessment of children with short stature for potential growth hormone deficiency. *IGF1* mutations lead to severe intrauterine growth restriction, postnatal growth failure, and developmental delay (Woods 1996 10), while *IGF2* mutations have been shown to be associated with Silver-Russell Syndrome (Begemann 2015 11). For IGF signalling to occur, it has been shown that IGFs must be released from IGF Binding Proteins (IGFBPs), which, among other functions, sequester IGFs and prolong their half life (David 2011 3, Bach 2018 12, Baxter 2000 13). Some IGFBPs are also necessary for localization and transport of IGFs. Specifically, IGFBP-3 controls IGF-II concentration (Ismayilnajadteymurabadi 2017 14), binding between 90-96% of free IGF-I and IGF-II (Jones 1995 15), and IGFBP-3 and IGF-II serum levels have been shown to be tightly correlated (r = .56) in adults (Probst-Hensch 2001 16). Measurement of intact IGFBPs are complicated due to their cleavage by the proteases, pregnancy-associated plasma protein A (PAPP-A) and A2 (PAPP-A2). IGFBP cleavage allows IGF release, binding to their cell surface receptor, IGF-1R, and downstream signalling. Multiple proteins in the IGF signalling pathway have been implicated by genome-wide association studies (GWAS) of both height and skeletal growth phenotypes (Wood 2014 17, Yengo 2018 18, Chan 2015 19); specifically, Wood et al reported that height GWAS identifies at loci or near *IGF1, IGF2, IGFBP3, IGFBP5, PAPPA*, and *PAPPA2*. GWAS of sitting height ratio, a measure of skeletal proportion, also implicate the *IGFBP3* locus. However, the mechanism by which the variants in these genes influence skeletal growth is still unknown. Further genetic study can offer insight into the interactions and shared underlying mechanisms of the GH system and human growth. Furthermore, genetic associations can enable causal inference and provide evidence regarding whether circulating levels of these proteins causally influence height and other skeletal growth phenotypes.

Previous work (Fujimoto 2020 20) described the distributions of serum concentrations for many of these proteins through childhood in a large cross-sectional cohort. In this report, we describe the distributions of IGF-II and IGFBP-5 levels and perform a GWAS of multiple circulating protein levels in participants from this same cohort to identify genetic variants that influence circulating protein levels during childhood (protein quantitative trait loci, pQTLs). We also explore the relationship between pQTLs for different growth-related proteins and skeletal growth phenotypes (including height and sitting height ratio), and identify surprising and distinct relationships between the genetic variants near genes encoding the growth-related proteins, the circulating protein levels, and the skeletal growth phenotypes.

## Results

### IGF-II and IGFBP-5 levels throughout childhood

Our group has previously reported distributions of total and free IGF-I, PAPP-A2, total and intact IGFBP-3 throughout childhood as well as the correlation between these proteins (Fujimoto 2020 20). Herein, we present additional data describing IGF-II and IGFBP-5 levels in the same cohort. Serum IGF-II levels increased in childhood with a subsequent flattening of the curves (Figure 1 A, B and Supplemental Table 3). Serum IGFBP-5 in females increased until approximately 13 years old and then flattened. Interestingly, serum IGFBP-5 levels in males remained flat until the age of 10-11 years and then progressively increased (Figure 1 C, D and Supplemental Table 4).

**Figure 1:**
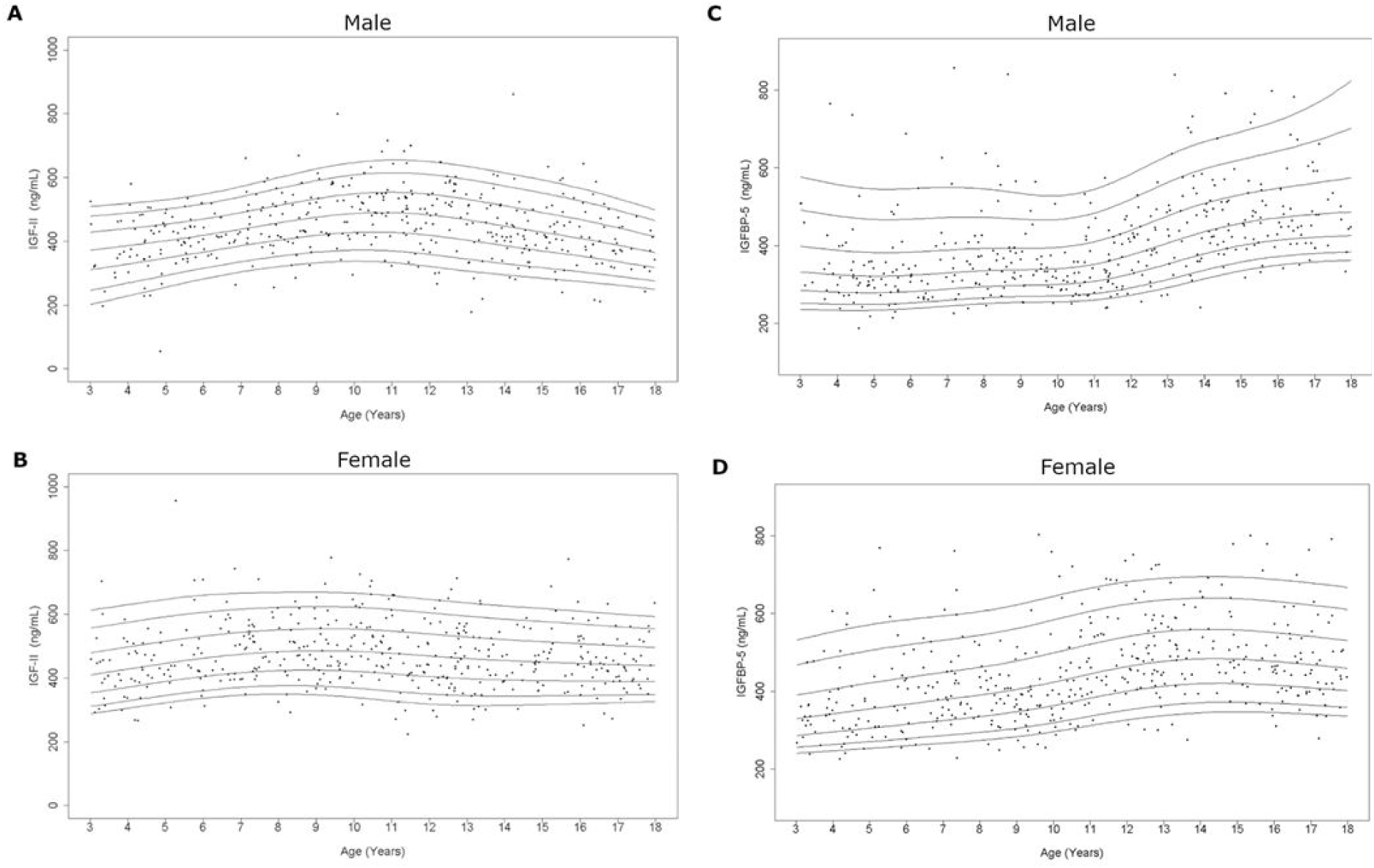
Serum IGF-II and IGFBP-5 curves in the study population. Age distribution of serum IGF-II and IGFBP-5 levels in males (A, C) and females (B, D) are represented respectively. The curves of the 5th, 10th, 25th, 50th, 75th, 90th, and 95th percentiles calculated by the BCT method are displayed.

Correlation analyses are presented in Table 1. IGF-II levels had a strong positive correlation with total IGFBP-3 in both sexes (M, F; r = 0.53, 0.51; P < 0.0001). However, IGF-II levels were weakly positively correlated with intact IGFBP-3 (r = 0.17, *P* = 0.0006) in males only. There were no significant correlations between IGF-II and height or BMI throughout childhood. There was no significant correlation between serum IGF-II and IGFBP-5 levels in either sex (Male, Female: r=-0.10 (*P* = 0.05), r=0.07 (*P*=0.13), respectively).

**Table 1:** Correlation analyses between IGF-II, IGFBP-5 and other variables and regression with Height and BMI.

| Variable |  | IGFBP-5 | Free IGF-I<sup>a</sup> | Total IGF-I<sup>a</sup> | Intact IGFBP-3 | Total IGFBP-3 | PAPP-A2<sup>a</sup> | Free / total IGF-I<sup>a</sup> | Height z-score | BMI z-score |
| --- | --- | --- | --- | --- | --- | --- | --- | --- | --- | --- |
| Males |  |  |  |  |  |  |  |  |  |  |
| IGF-II<br>(N = 405) | r | -0.10 | -0.15 | 0.07 | 0.17 | 0.53 | 0.02 | -0.19 | -0.015 | -0.007 |
|  | P value | (0.05) | (0.003) | (0.17) | (0.0006) | (<0.0001) | (0.65) | (0.0002) | (0.77) | (0.89) |
| IGFBP-5<br>(N = 405) | r |  | 0.39 | 0.44 | 0.32 | 0.008 | -0.14 | -0.009 | 0.12 | -0.02 |
|  | P value |  | (<0.0001) | (<0.0001) | (<0.0001) | (0.87) | (0.004) | (0.86) | (0.02) | (0.71) |
| Females |  |  |  |  |  |  |  |  |  |  |
| IGF-II<br>(N = 431) | r | 0.07 | 0.06 | -0.05 | 0.04 | 0.51 | 0.12 | 0.10 | -0.028 | -0.045 |
|  | P value | (0.13) | (0.22) | (0.27) | (0.57) | (<0.0001) | (0.02) | (0.03) | (0.56) | (0.35) |
| IGFBP-5<br>(N = 431) | r |  | 0.41 | 0.51 | 0.32 | 0.18 | -0.10 | -0.04 | 0.04 | -0.029 |
|  | P value |  | (<0.0001) | (<0.0001) | (<0.0001) | (0.0001) | (0.04) | (0.37) | (0.37) | (0.55) |

a The natural log-transformation was used for statistical analysis of free and total IGF-I, PAPP-A2 and free / total IGF-I ratio.
Note: Bivariate Pearson correlation analyses were performed. The r means the Pearson correlation coefficient. A P-value less than 0.05 was considered statistically significant.

IGFBP-5 levels had a positive correlation with free and total IGF-I in both sexes (M, F; free IGF-I, r = 0.39, 0.41; total IGF-I, r = 0.44, 0.51; all P < 0.0001). IGFBP-5 levels were significantly correlated with intact IGFBP-3 (r = 0.32; P < 0.0001) in both sexes. There were no significant correlations between IGFBP-5 and height or BMI throughout childhood.

### Genetic analyses

#### Genome-wide association studies for protein levels

GWAS was performed for each of the 7 phenotypes (Free IGF-I, Total IGF-I, IGF-II, IGFBP-5, Total IGFBP-3, Intact IGFBP-3, PAPP-A2) as well as two derived ratio phenotypes (Free / Total IGF-I, Intact IGFBP-3 / Total IGFBP-3) and two conditional phenotypes (IGF-II conditioned on Intact IGFBP-3, IGF-II conditioned on Total IGFBP-3). This was done for two populations (African American and European American) for each sex strata (sex-combined, male, female), while adjusting for covariates described in the methods section. Directly genotyped or imputed single nucleotide polymorphisms (SNPs) were analyzed in the GWAS. Fixed-effects meta analysis was performed using METAL to combine GWAS results across the two populations, and likely independent SNPs were selected as having the best association at each 1 Mb genomic locus (“lead SNPs”). In total, we identified 17 independent lead SNPs with association P values below a commonly used genome-wide significance threshold of 5×10^−8^ (Supplemental table 1). Strongest associations were identified at the rs2854746 SNP within the *IGFBP3* locus, associated with both IGF-II and Total IGFBP-3 levels, and the rs11575194 SNP within the *IGFBP5* locus, associated with IGFBP-5 levels. The rs2854746 variant is a previously known expression QTL (eQTL) and protein QTL (pQTL) (Kaplan 2011 21, lead snp R^2^=0.883 Machiela 2015 46) (also reported in Deal 2001 22, Canzian 2006 23, Schumacher 2010 24, Evans 2014 25, and Teumer 2016 26), associated with levels of *IGFBP3* expression and IGFBP-3 in serum but has not to our knowledge been previously associated with IGF-II levels. By conditioning on IGFBP-3 levels, we show the novel rs2854746 association with IGF-II is independent of intact IGFBP-3 and is attenuated but remains significant when conditioning on total IGFBP-3 (Supplemental Figure 2). The rs11575194 variant is a known pQTL for OSBPL-11 (gtex 27, Sun 2018 28) but has not to our knowledge been reported to be associated with IGFBP-5 levels. 17 additional associations reached the genome-wide significance threshold, across a range of phenotypes as described in Supplementary table 1.

#### Overlap with findings from GWAS of skeletal growth phenotypes

GWAS was performed for standing height (referred to simply as “height”), SHR (sitting height to standing height ratio), and birth weight using data from the UKBiobank (see Methods). The rs2854746 SNP, which is associated with total IGFBP-3 and IGF-II levels, lies near the *IGFBP3* gene and is also significantly associated with both height (*p* = 5.0×10^−12^) and SHR (*p* = 3.8×10^−69^), but not with birth weight. However, although rs2854746 was also the strongest associated SNP for SHR in the region, it was only weakly correlated (*R*^2^ = 0.0045, 0.0001 respectively) with the 2 lead height SNPs in the region (rs11455214, *p* = 1.7×10^−25^, rs723149, *p* = 3.2×10^−50^), consistent with earlier suggestions that these might reflect different signals at the same locus (Chan 2015 19). The rs11575194 SNP, which is associated with IGFBP-5 levels, lies near the *IGFBP5* gene, and is associated with birth weight (*p* = 6. ×10^−9^) but not with height or SHR, despite the fact that other SNPs near *IGFBP5* are strongly associated with height (lead SNP is rs2241192, *p* = 1.3×10^−14^, *R*^2^ = 0.0004, rs1478575, *p* = 8.9×10^−58^, *R*^2^ = 0 in 1000 Genomes).

To look for other phenotypes that could be influenced by the levels of IGF-related proteins, we also performed an association study across many other phenotypes (pheWAS) using the T2Dportal (T2D portal 29,30), the UKB pheWEB (pheweb http://pheweb.sph.umich.edu/) (Supplemental Figure 3), and the GWAS catalogue (Buniello 2019 31) for the lead SNPs affecting total IGFBP-3 and IGFBP-5 levels. Although IGF-related proteins have been suggested as risk factors for cancer, neither lead SNP has been identified in the GWAS catalog as being strongly associated with colorectal, breast, or prostate cancers. The only associations surviving multiple testing correction (besides skeletal growth phenotypes) were between rs2854746 in *IGFBP3* and multiple measures of bioelectrical impedance (legs, whole body, and arms), as well as diastolic and systolic blood pressures.

#### Conditional analyses to distinguish association signals at the IGFBP3 *and* IGFBP5 *loci*

Although the weak correlations between lead SNPs for height and total IGFBP-3 levels suggest that these associations are distinct, the presence of multiple signals of association at a locus can complicate interpretation of simple correlation measures between lead SNPs. To more formally evaluate whether the associations at the *IGFBP3* locus with height and total IGFBP-3 levels actually reflect distinct associations, and hence distinct mechanisms, we performed serial conditional analyses. For each phenotype, we performed association analysis for the SNPs across the locus, conditioning on the lead SNP for that phenotype, and then continued to condition on the strongest remaining SNP associated with that phenotype until no genome-wide significant signal of association remained at the locus (see Methods). When conditioning in turn on each of 2 total IGFBP-3 associated independent SNPs in the *IGFBP3* locus, we observed minimal change in the strength of the height associations in the region. Similarly, when conditioning in turn on 13 height-associated SNPs, we observe that associations with total IGFBP-3 protein levels remained robust (Figure 2a, Supplemental Figure 4a-b). In contrast, when we performed similar analyses comparing associations of total IGFBP-3 levels and SHR, the signals associated with SHR were dramatically attenuated when conditioning on the two total IGFBP-3 associated SNPs, and, comparably, total IGFBP-3 association dramatically attenuated when conditioning on the 6 SHR associated SNPs (Figure 2b, Supplemental Figure 4c-d). Together, these analyses indicate that the associations with height at the *IGFBP3* locus are independent of associations with serum total IGFBP-3 protein levels but the effects on SHR and total IGFBP-3 levels likely represent the same signal.

**Figure 2:**
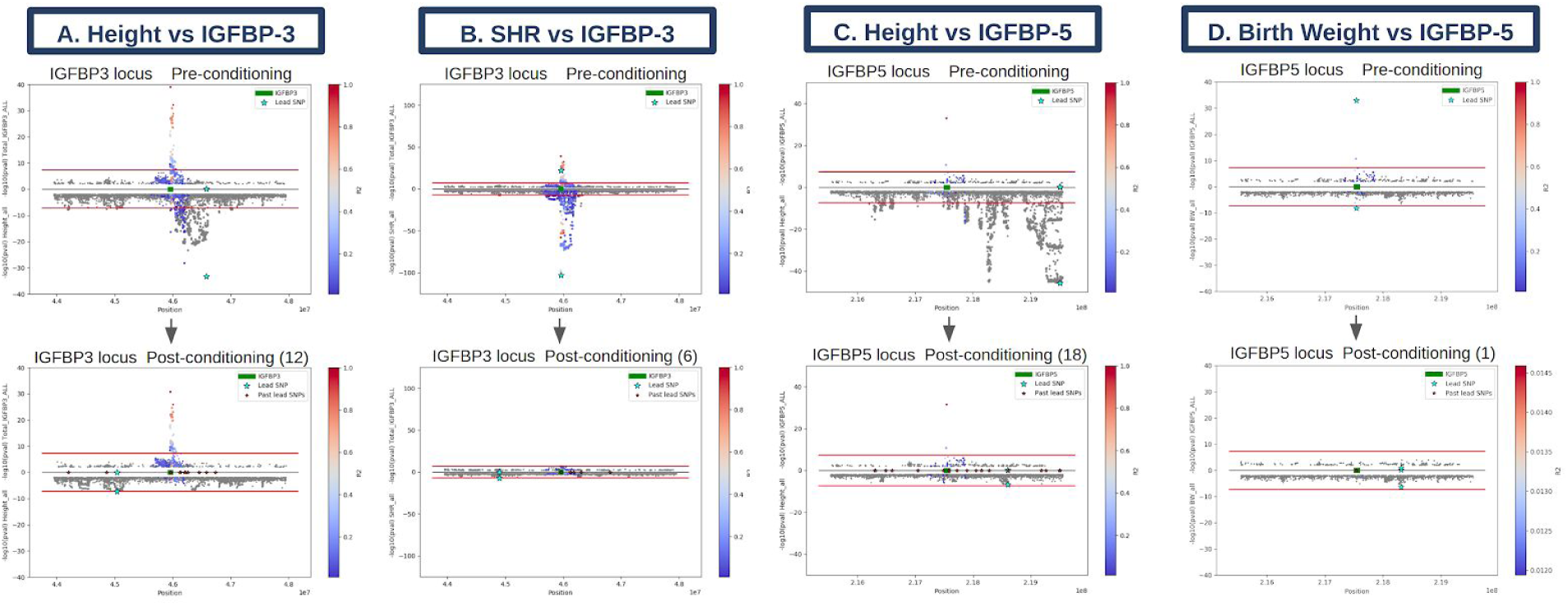
Conditional analysis. Serial conditional analyses were performed comparing **A**. total IGFBP-3 association signal with height associations in the region. After conditioning on 13 nearby (2MB window) height lead SNPs, SNPs in the region remain associated with total IGFBP-3 levels. **B**. total IGFBP-3 with SHR. After conditioning on 6 nearby SHR lead SNPs, SNP association with total IGFBP-3 levels is attenuated. **C**. IGFBP-5 with height. After conditioning on 18 nearby height lead SNPs, SNPs in the region remain associated with IGFBP-5 levels. **B**. IGFBP-5 with birth weight. After conditioning on the 1 nearby BW lead SNP, SNP association with IGFBP-5 levels is attenuated.

We performed similar serial conditional analyses at the *IGFBP5* locus to examine the relationship between the associations with IGFBP-5 levels, height, and birth weight. The signal of association with IGFBP-5 is vastly attenuated when conditioning birth weight signals in the region (and vice versa), suggesting that these reflect the same association signal (Figure 2d, Supplemental Figure 4f). In contrast, when we serially conditioned on 18 height signals in the region, we observed minimal attenuation of the IGFBP-5 associations, and conditioning on IGFBP-5 associated SNPs did not substantially attenuate the height signals (Figure 2c, Supplemental Figure 4e). Thus, similar to our observations at the *IGFBP3* locus, these results indicate that associations with height are distinct from the associations with serum levels of IGFBP-5, but the serum levels are associated with a distinct skeletal growth phenotype (in this case, birth weight).

Genetic studies of height have identified signals near other genes encoding IGF-related proteins (Wood 2014 17, Yengo 2018 18, Marouli 2017 32). To further explore the relationship between serum levels of IGF-related proteins and height association signals near the genes encoding these proteins, we focused on the height-associated SNPs in the loci containing the *IGF1, IGFBP3, PAPPA2, IGF2*, and *IGFBP5* genes. We tested nine common genome-wide significant height SNPs for association with protein levels. Results of each SNP association are presented in Supplementary Table 2. The only significant association (after correction for multiple comparisons) was between the rs1044299 SNP at the *PAPPA2* locus and serum PAPP-A2 levels (*p* = 6.85×10^−5^). Conditioning on one of the height signals near *PAPPA2* (rs1040457) vastly attenuated this association with PAPP-A2 serum levels; however, the most significant height lead SNP (rs10913200) is independent of the association with PAPP-A2 levels (Supplemental Figure 5). Thus, as with *IGFBP3* and *IGFBP5* the effects on height of variants at the other IGF-related genes are not mediated by serum levels, with the possible exception of one variant near *PAPPA2*.

#### Mendelian Randomization

To evaluate causal relationships between identified associations at *IGFBP3* and *IGFBP5* and skeletal growth phenotypes, we performed Mendelian Randomization. In this method, a SNP associated with the proposed causal variable is used as an instrumental variable (IV) and the IV’s association is tested with the outcome variable. We first used rs2854746 as an instrumental variable (IV) for total IGFBP-3 levels (after conditioning on other height signals in the region). We show increasing total IGFBP-3 levels by 1 standard deviation (SD) is associated with a 0.061 SD increase in SHR (95% CI: [0.053, 0.069]) and a much smaller 0.010 SD decrease in height (95% CI [-0.018, -0.0023]) (Figure 3A). Similarly, using rs11575194 as the IV for IGFBP-5 levels (after conditioning on other height signals in the region), we show increasing IGFBP-5 levels by 1 SD is associated with a 0.027 SD decrease in birth weight (95% CI [-0.036, -0.018]) and a smaller 0.010 SD decrease in height (95% CI [-0.017, -0.004]); the IGFBP-5 - birth weight relationship may act through correlated measures such as fetal serum levels. These results suggest that the main detectable causal effects of variation in serum levels of total IGFBP-3 and IGFBP-5 in childhood are on skeletal proportion (SHR) and birth weight, respectively.

**Figure 3:**
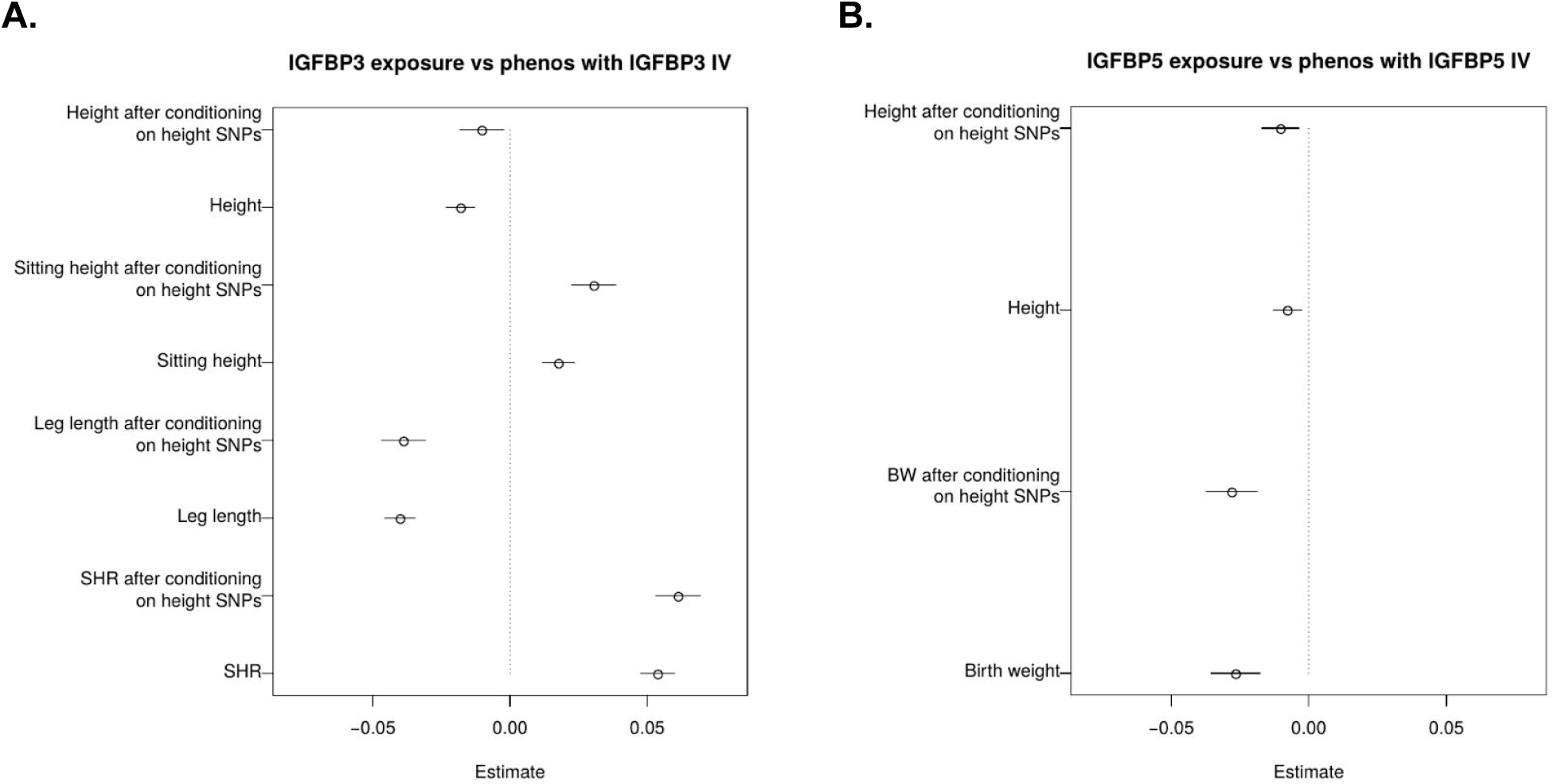
MR. Mendelian randomization was performed; rs2854746 was used as an instrumental variable (IV) for total IGFBP-3 levels and rs11575194 was used as an IV for IGFBP-5 levels. GWAS was performed conditioning on nearby height SNPs to ameliorate their effect on effect size estimates. **A**. MR analyses show change in total IGFBP-3 levels correlate to more standard deviation changes in SHR than in height. **B**. MR analyses show change in IGFBP-5 levels correlate to more standard deviation changes in birth weight than in height.

**Figure 4:**
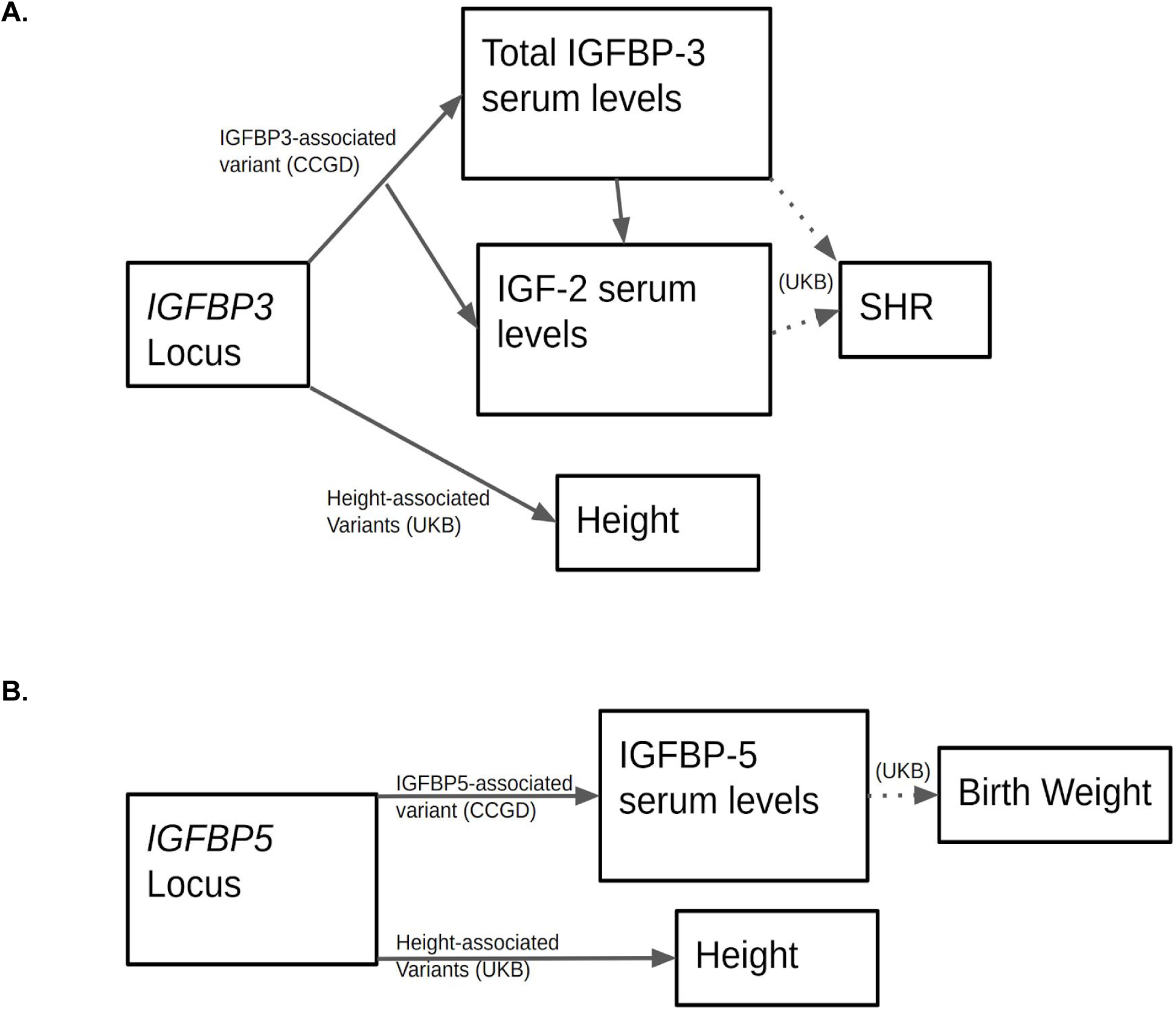
Schematic. **A.** The *IGFBP3* locus is associated with both height and SHR. The total IGFBP-3 association, with lead SNP rs2854746, colocalizes with the SHR association but not the height association. As such, it is possible total IGFBP-3 serum levels and possibly IGF-II serum levels modulate SHR but not height, and other regulatory mechanisms of *IGFBP3* in other tissues or other time points are responsible for *IGFBP3*’s association with height. Dashed lines mark interactions for which there is most evidence in MR analyses. **B**. Similarly, the *IGFBP5* locus is associated with both height and birth weight. The IGFBP-5 association, with lead SNP rs11575194, colocalizes with the birth weight association. Therefore, it is possible the *IGFBP5* locus affects birth weight through IGFBP-5 serum levels, and interacts with height not through serum levels, but rather in some other tissues or time points.

## Discussion

The GH and IGF regulatory system is critical for understanding biology driving human growth, height, and related phenotypes. Recent work has explored IGF protein interactions and their regulation of height-relevant biology, across multiple age ranges (Fujimoto 2020 20, Kloverpris 2013 33, Steinbrecher 2017 34). In this study, we performed GWAS of measured serum levels of IGF-related proteins throughout childhood, and identified 17 associated genetic loci. These included strong effects at the *IGFBP3* locus on total IGFBP-3 and IGF-II levels and at the *IGFBP5* locus on IGFBP-5 levels. We used these associations to explore the relationship between measured serum levels with mechanisms driving height and related phenotypes, and provided evidence that serum levels of IGF-II, total IGFBP-3, and IGFBP-5 do not appear to be causal for height but instead have evidence for causality for other skeletal growth phenotypes, notably total IGFBP-3 and/or IGF-II with sitting height ratio and IGFBP-5 with birth weight.

Normal levels of GH-IGF axis-related proteins are well described in adults and through childhood. Large population based studies have examined the concentrations of serum total IGF-I and total IGFBP-3 throughout the lifespan (Friedrich 2014 35, Bidlingmaier 2014 36). Our group has measured serum levels of PAPP-A2 and GH-IGF axis-related proteins (now including IGF-II and IGFBP-5) throughout childhood, and performed correlation analyses of all the analytes to investigate the interplay between the various proteins.

IGF-related proteins are known to play a role in regulation of growth and are canonically understood to affect human height; as such, changes in levels or function would logically be assumed to affect overall growth and therefore adult height. Because genetic factors affect both protein levels and measures of human growth, genetic studies can be used to gain insights into the relationships between serum protein levels and skeletal growth. In the GWAS performed in this study, we confirm prior observations that there are signal(s) of association with height as well as with other skeletal growth phenotypes near the *IGFBP3* locus. Surprisingly, the variant associated with serum levels of total IGFBP-3 and IGF-II is far more strongly associated with adult SHR than with adult height; indeed, as suggested by earlier studies (Chan 2015 19) our conditional analyses indicate that while SHR and protein levels share a common association at the *IGFBP3* locus, the associations with height and total IGFBP-3/IGF-II protein levels are independent of each other. These findings support a surprising conclusion, wherein serum levels of total IGFBP-3, IGF-II, or other unmeasured correlated variables causally affect predominantly SHR, rather than height. It follows that variation in human height is not caused by total IGFBP-3 nor IGF-II levels, as measured in the serum. Further, because there are at least 13 variants clustered near the *IGFBP3* gene that are associated with height but not with total IGFBP-3 levels, our results suggest that regulation of expression of *IGFBP3* somewhere in the body does influence overall height, but that *IGFBP3* expression in this as yet undefined place (or time) in the body does not correlate with measured serum levels of total IGFBP-3 in childhood. Future examination of the regulatory elements near the *IGFBP3* variants associated with height should provide clues as to the time(s) and place(s) where total IGFBP-3 influences overall growth.

It is also curious that the variant at *IGFBP3* that is associated with serum levels affects SHR but not overall height - although many variants affect both SHR and height, some, like this *IGFBP3* variant, only have a detectable effect on SHR. Because overall height is composed of sitting height (head+spine) and leg length, the effect on SHR without an effect on overall height means that the variant must have symmetric and opposing effects on head+spine and leg length. We are not aware of an obvious mechanism for such coordinated and opposing effects on skeletal growth, although one could postulate a mechanism that somehow regulates overall height, meaning that an effect on one component of height (for example on leg length) could cause an indirect compensation reflected in the other component of height (for example, head+spine). Alternatively, total IGFBP-3 or IGF-II serum levels (or something correlated with these) could have two independent and opposing effects on head+spine and leg length that just happen to cancel each other out in their effect on total height.

Interestingly, the variant associated with total IGFBP-3 levels is also known to influence expression of *IGFBP3* mRNA in multiple tissues (gtex 27). However, these tissues (Heart - Left Ventricle, Thyroid, Muscle - Skeletal, Lung, Brain - Cerebellum, Artery - Aorta, Artery - Tibial, Esophagus - Muscularis https://gtexportal.org/home/snp/rs2854746 27) do not include the liver, which is presumed to be the main source of circulating IGFBP-3. Furthermore, this variant is associated only with total circulating IGFBP-3 but not with intact IGFBP-3 levels, which are thought to reflect the biologically active form of IGFBP-3. Because this variant influences SHR, these observations collectively raise the possibility that non-intact forms of IGFBP-3 may affect skeletal proportion either directly or indirectly (for example through IGF-II levels or other correlated measures) and that the relevant expression of IGFBP-3 is not mediated through changes in *IGFBP3* expression in the liver.

We also observed variation at the *IGFBP5* gene locus that is associated with IGFBP-5 levels. Like *IGFBP3*, the *IGFBP5* locus is known to be associated with human height (Wood 2014 17, Yengo 2018 18). However, as we observed for the *IGFBP3* locus, the associations with height and IGFBP-5 protein levels are independent, as shown by conditional analysis; this indicates circulating serum levels of IGFBP-5 are likely not mediating the effect of IGFBP-5 on height. In contrast to variation at the *IGFBP3* locus, the variation at *IGFBP5* that influences protein levels is not associated with SHR; however, it is associated with birth weight, suggesting that serum IGFBP-5 levels are important for prenatal growth through a mechanism that is distinct from the influence of IGFBP-5 on adult height. As with *IGFBP3*, the influence of variation in *IGFBP5* expression on height must occur at a time and/or place that is not reflected in circulating protein levels in childhood.

We also performed an analysis of the lead height-associated SNPs near the remaining genes encoding IGF-related proteins in our study. None of the lead height SNPs were significantly associated with serum levels of the proteins, with the exception of one variant near *PAPPA2*. Again, this suggests that serum levels of these proteins during childhood are not necessarily determinative of adult height. These results do not mean that these proteins can not serve as clinical biomarkers but it does imply the existence of mechanisms for these genetic associations with height that are not reflected in measured serum proteins.

Beyond the signals at *IGFBP3* and *IGFBP5*, 17 additional genome-wide associations were identified with IGF-related protein levels. None had clear colocalization with either the genes encoding these proteins or other skeletal growth phenotypes, but may provide clues as to the regulation of these proteins. To allow further exploration of these results, we will provide access to a genome-wide set of summary association statistics.

This study has a number of limitations. First, the sample size for serum protein associations is small for a GWAS study, although the presence of highly significant associations with large effects on measured serum levels (as has been seen in other studies) shows that we had adequate power to successfully identify these strong associations. Larger sample sizes would also allow for additional analyses, such as estimates of heritabilities and genetic correlations for serum protein levels, as well as identification of variants with more modest effects. Second, we provide evidence that some association signals overlap (such as total IGFBP-3 serum levels and sitting height ratio), but methods to rigorously evaluate colocalization (COLOC 37) or genetic correlation (ldsc 38) are limited in small samples and these analyses are further complicated by the presence of many independent associations in each region. We also did not have parental genotypes, so could not directly analyze parent-of-origin effects, which decreases power at imprinted loci (for example, the *IGF2* gene is imprinted).

The GWAS analysis was corrected for population structure in the data using principal components of genetic ancestry, and our sample was stratified by ancestry before meta-analysis, but there may still be confounding from residual population stratification reflected in the summary statistics. In addition, data from this study were largely derived from European descent individuals (both in the UKB and Cincinnati cohorts), limiting the generalizability to non-European ancestry; it would be beneficial to expand these analyses of serum protein levels to other ancestries.

Because the rs2854746 in *IGFBP3* is a missense variant, it is theoretically possible that this variant affects binding of the ELISA probe, and is not actually associated with circulating serum protein levels. However, the amino acid altered by rs2854746 does not overlap the probe binding location, and the association with IGF-II levels is strongly indicative of a true biological effect, rather than a laboratory artifact. As with all GWAS associations, functional follow-up of this and the other loci we identified will be necessary to elucidate the specific mechanism of action of these findings.

In conclusion, this study identifies strong genetic associations with measured serum levels of IGF-related proteins, particularly between IGF-II and total IGFBP-3 serum protein levels and variation at the *IGFBP3* locus, and between IGFBP-5 protein levels and variation at the *IGFBP5* locus. These associations are distinct from and independent of the associations with adult height that have been observed at these loci. Rather, the *IGFBP3* variant rs2854746 is strongly associated with SHR, and the *IGFBP5* variant rs11575194 is associated with birth weight. These findings suggest that genetically-encoded variation in *IGFBP3* and *IGFBP5* expression affects height in ways that are as yet unknown but are independent of measured serum levels during childhood (for example, local paracrine effects in the growth plate or developing skeleton, or effects on serum levels in fetal life or infancy). By contrast, effects on skeletal proportion and birth weight may be more directly mediated through measured serum levels of total IGFBP-3 and IGFBP-5, respectively (and/or by correlated phenotypes, such as IGF-II levels). Further functional studies are needed to elucidate the mechanism of effect of variants in these genes on height, skeletal proportion, and birth weight, and specifically whether and how their effects are mediated by local or systemic changes in the level or activity of growth factor-related proteins, possibly at specific times in development. Taken together, these findings challenge our understanding of how well-known growth factors affect skeletal growth and proportion in children.

## Materials and Methods

### Study population and design of the Cincinnati Genomic Control Cohort

Subject selection for this cohort has been previously described (Fujimoto 2020 20). The study subjects and data were derived from the Cincinnati Genomic Control Cohort (CGCC), which is a community-based cohort comprised of 1,020 -children; 838 were included after filtering for obesity, undernutrition, or severe short or tall stature. The study was reviewed and approved by the institutional review board at Cincinnati Children’s Hospital Medical Center.

### Anthropometric data and samples

All participants had their height and weight measured by a single investigator as previously described (Fujimoto 2020 20). The datasets of CDC 2000 growth charts were used to calculate age- and sex-adjusted Z-scores (Kuczmarski RJ 2002 39). Blood samples were taken from the participants to obtain serum and genomic DNA at once. Serum samples were isolated with a standard coagulation and centrifuge protocol and frozen at −80°C within 2 hours of collection (Fujimoto 2020 20). The serum samples were stored at −80 ° C for more than 8 years without thawing.

### IGF-related Protein Measurements

Serum concentrations of IGF-II, IGFBP-5, free and total IGF-I, intact and total IGFBP-3, and PAPP-A2 were measured using a commercially available enzyme-linked immunosorbent assays (ELISA) kits (AL-121, 122, 149, 120, 109, 131 and 127; Ansh labs, TX, USA) as previously published (DiPrisco 2019 40, Fujimoto 2020 20). The limits of detection and limits of quantification, imprecision, linearity, and recovery were determined for each assay. The IGF-II assay demonstrates no significant (< 0.1 %) cross-reactivity with IGFBP-2, -3, -4, -5, and IGF-I to 1000 ng/mL. The IGFBP-5 assay demonstrates no significant (< 0.1 %) cross-reactivity with IGFBP-2, -3, -4, IGF-I, and IGF-II to 1000 ng/mL.

### Study-based IGF-II and IGFBP-5 curves generation

Age- and sex-specific IGF-II and IGFBP-5 curves in this study were created using the R package as previously published (Fujimoto 2020 20). Curve development was done using the Box-Cox t (BCT) distribution. The models included the median, a coefficient of variation, and a measure of skewness and/or kurtosis. The study based IGF-II and IGFBP-5 curves were constructed with the percentile lines of 5, 10, 25, 50, 75, 90, and 95^th^ for each analyte (Figure 1). The percentile data for each of these analytes is presented in Supplemental Tables 3-4.

### GWAS data description

#### Genotyping and quality control of genotype data

The single nucleotide polymorphism (SNP) genotyping was carried out previously at Cincinnati Children’s Hospital Medical Center (CCHMC) using two different arrays (Affymetrix™ or Omni-5™ platforms).

Ancestry was estimated by principal components (PC) analysis: the genotype data for each participant was projected onto PCs from the 1000Genomes samples (1kgP3 paper 2015 41) to define two groups of participants based on their estimated continental ancestries; participants were placed into groups whose genotype data were broadly consistent with either European ancestry or with admixed African and European ancestry. Participants were removed based on sex missingness or mismatch, genotype call rate (< 95%), and haplotype missingness (*p* > 1×10^−8^). Variants were filtered using Plink2-1.90b3.32 (Plink Chang 2015 42) based on multiple criteria: monomorphic sites, site missingness (>10%), heterozygosity (+/- 4 sd), hardy-weinberg equilibrium (*p* > 1×10^−7^). Genotype data was aligned to 1000Genomes phase 3 data (1kgP3 paper 2015 41) and imputation was performed using the University of Michigan imputation server (Das 2016 43). Variants were filtered out based on low imputation info score (R^2^ < 0.3) and minor allele frequency (<0.01).

#### Phenotyping

Height z-scores and BMI z-scores were generated using the 2000 CDC growth charts. Subjects were excluded for extreme height or BMI z-scores (z > 3 or z < -3), chronic illness, and pregnancy.

#### GWAS analysis of Cincinnati and UKB cohorts

GWAS in the Cincinnati cohort was performed on samples stratified by estimated ancestry (European ancestry or African-American ancestry) and by sex (male only, female only, and all). Prior to running the GWAS, derived traits were calculated, and adjusted phenotype values were adjusted PC1-10, and sex (for the all analysis); for conditional traits (IGF-II conditioned on Intact IGFBP-3 and IGF-II conditioned on Total IGFBP-3), residuals were calculated by conditioning the IGF-II phenotype on respective IGFBP-3 levels. Inverse normal (rank-based) transformation was performed on the residuals after these adjustments. GWAS was performed using linear regression with plink software. Cross population analyses were fixed-effects meta analyzed using METAL (Willer 2010 44). Independent loci were selected by identifying lead SNPs in 1 Mb windows.

GWAS was performed in the UK Biobank for locus comparison for traits height and sitting height ratio. Analyses were performed on individuals of European ancestry (N=455146) and stratified by sex. Phenotype values were adjusted for age and sex (in the unstratified analysis). Residuals were filtered for extreme outliers (+/- 4 SD) and inverse normal transformation was performed. GWAS was performed using Bolt-lmm v2.3 (Loh 2019 45), adjusting for array type and 10 PCs of ancestry.

#### Serial Conditioning for eQTLs

For the *IGFBP3* locus, serial conditional analyses were performed to evaluate independence with nearby height associations. For the first analysis, GWAS was performed conditioning on the height lead SNP (within +/- 2MB). This was repeated, conditioning on the first height lead SNP and additionally next height lead SNP, and iterated; eventually after 13 steps, GWAS was performed conditioning on 13 height lead SNPs (stopping when no genome-wide significant associations p < 5×10^−8^ remained). At the same locus, this analysis was repeated conditioning on IGFBP-3 lead SNPs (2 iterations, 2 IGFBP-3 lead SNPs). Similar conditional analyses were repeated for IGFBP-3 and SHR, IGFBP-5 and Birth weight, and IGFBP-5 and height. Conditional analyses were also performed for PAPP-A2 and height, with the modification that, when conditioning on PAPP-A2 associated SNPs, SNPs are considered lead SNPs if GWAS p < 1×10^−4^ (rather than 5×10^−8^, to investigate weak associations in the region).

#### Mendelian Randomization analyses

We performed Mendelian Randomization to evaluate the causal relationship between protein levels and relevant phenotypes. For each pair of traits (IGFBP-3 levels with each of height, SHR, sitting height, and leg length; IGFBP-5 levels with each of height and BW), effect sizes were estimated using the Inverse-variance weighted (IVW) model. For pairs including IGBFP-3 levels, rs2854746 was used as the instrumental variable; for trait pairs involving IGFBP-5 levels, rs11575194 was used. Effect sizes for protein level associations were estimated in GWAS in CCHMC described above; effect sizes for anthropometric phenotypes were estimated in GWAS of UKB as described above. To avoid inflation of effect estimates by nearby but distinct height associated variants, each analysis was repeated using GWAS conditioned on all GWS height signals in each locus (13 SNPs for *IGFBP3*, 18 SNPs for *IGFBP5*). MR was performed using the MendelianRandomization 0.3.0 R package (https://cran.r-project.org/web/packages/MendelianRandomization/vignettes/Vignette_MR.pdf)

### Statistical Analysis for Association between the Dependent Variables

As previously described (Fujimoto 2020 20), data were imported into SAS®, version 9.4 (SAS Institute, Cary NC), for data management and analysis except for GWAS analysis. Continuous data were examined for distributional properties and potential outlying values. After outliers removed, three of the dependent variables, free IGF-I, PAPP-A2, and total IGF-I were transformed using the natural logarithm for analysis. General linear models were used to examine changes over age for the dependent variables and differences by sex and the interaction of sex and age. Pearson correlation was used to examine the association between the dependent variables. A P-value of <0.05 was considered statistically significant.

## Acknowledgements

The authors thank Lisa Martin and Susan Thompson and the entire Cincinnati Genomics Control Cohort team for providing the samples for the study. The authors also thank Melissa Andrew and Leah Tyzinski for their assistance with sample handling. Furthermore, the authors thank Joanne Cole for her help guiding GWAS analyses, Christina Astley for support with MR, and Yu-han Hsu, Rebecca Fine, and Tiffany Amariuta for many useful discussions.

---

Supplemental tables exist as excel (uploaded) or in google drive.

Supplemental Table 1: Lead SNPs for 8 serum protein level GWASs https://docs.google.com/spreadsheets/d/1vF3y_dXn1B2tkzXhLWnhS3KSBEs0xaZo3c6pY0Kp-uA/edit#gid=923308025

Legend:

24 lead snps were identified in GWAS of 7 protein level phenotypes, two derived ratio phenotypes, and two conditional phenotypes in male-female- or all-analyses. The 2 most robust associations (identified in male-, female-, and all-analyses) were near *IGFBP3* and *IGFBP5*.

Supplemental Table 2: Protein level associations with Height SNPs near genes https://docs.google.com/spreadsheets/d/1vF3y_dXn1B2tkzXhLWnhS3KSBEs0xaZo3c6pY0Kp-uA/edit#gid=359130661

Legend:

Nearby height lead SNPs for each protein studied are reported, including the protein level association p-value for said SNP. Most are not associated, but the *PAPP-A2* loci’s rs1044299 is significantly associated with PAPP-A2 serum levels (although not genome-wide associated). These findings suggest height associated SNPs for all except maybe PAPP-A2 do not act through serum protein levels.

Supplementary Table 3: The percentile table of IGF-II. https://docs.google.com/spreadsheets/d/1vF3y_dXn1B2tkzXhLWnhS3KSBEs0xaZo3c6pY0Kp-uA/edit#gid=1596592799

Legend: The estimated percentile values for IGF-II serum levels in males and females at the every age in our dataset are calculated by the Box-Cox t (BCT) method which is an extended lambda-mu-sigma (LMS) method to estimate the median, coefficient of variation, and skewness.

Supplementary Table 4: The percentile table of IGFBP-5. https://docs.google.com/spreadsheets/d/1vF3y_dXn1B2tkzXhLWnhS3KSBEs0xaZo3c6pY0Kp-uA/edit#gid=1270265751

Legend: The estimated percentile values for IGFBP-5 serum levels in males and females at the every age in our dataset are calculated by the Box-Cox t (BCT) method which is an extended lambda-mu-sigma (LMS) method to estimate the median, coefficient of variation, and skewness.

**Supplemental figure 1:**
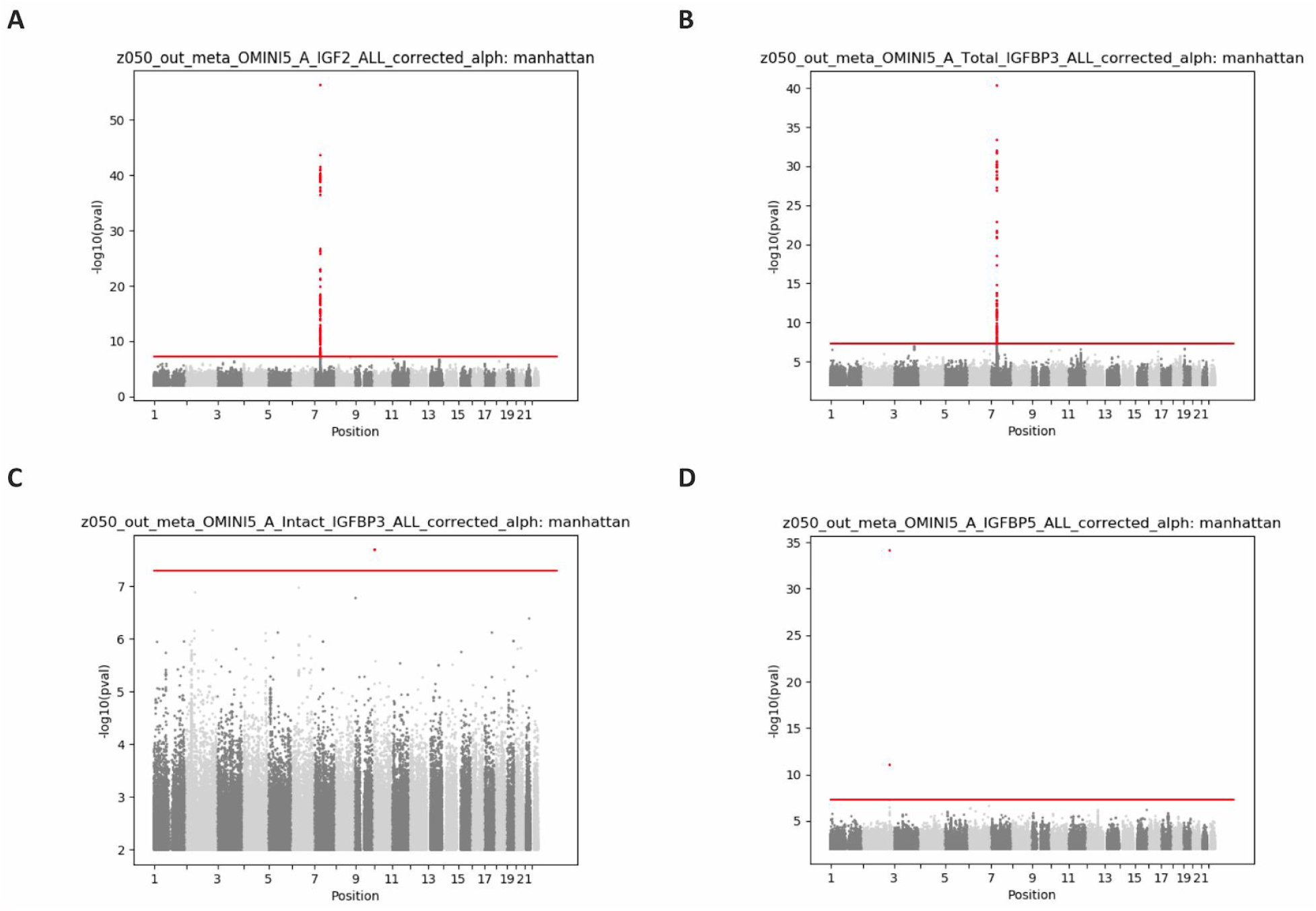
GWAS of A Serum IGF-II, B Total Serum IGFBP3, C Intact Serum IGFBP3, D Serum IGFBP5. GWAS meta-analysis associations for select serum phenotypes. **A**. IGF-II serum levels are genome wide significantly (gws) associated with 1 locus, occurring near *IGFBP3*. The lead snp is rs2854746. **B**. Total IGFBP-3 serum levels are gws associated with 1 locus near *IGFBP3*. The lead snp is rs2854746. **C**. Intact IGFBP-3 serum levels are not associated with any locus near *IGFBP3*, but are associated with a locus on chromosome 10. The nearest gene is *CALML3*. **D**. IGFBP-5 levels are gws associated with 1 locus near *IGFBP-5*. The lead snp is rs11575194.

**Supplemental figure 2:**
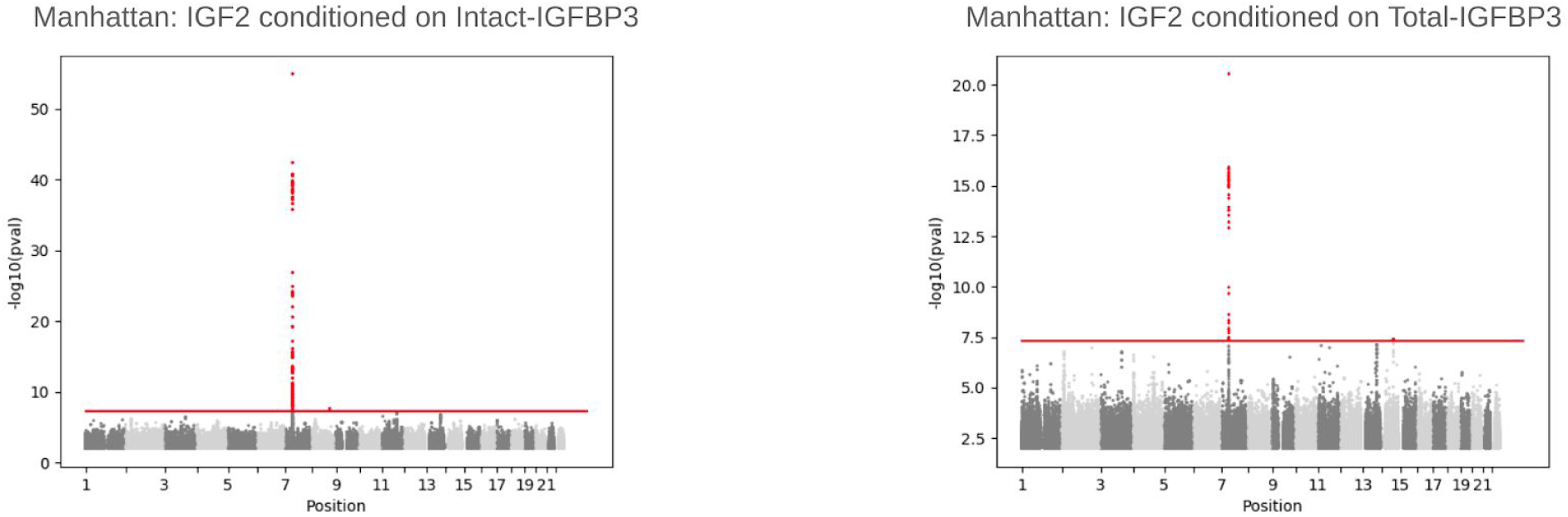
IGF-II association conditioned on intact- and total-IGFBP-3. Genome-wide associations from meta-analysis (A) with IGF-II conditioned on Intact-IGFBP3 and (B) with IGF-II conditioned on Intact-IGFBP3. Red line marks p=5e-8. Conditioning on intact IGFBP-3 levels minimally affects the observed association; conditioning on total IGFBP-3 attenuates the association with the *IGFBP3* locus. Both analyses remain significantly associated *IGFBP3* locus

**Supplemental figure 3:**
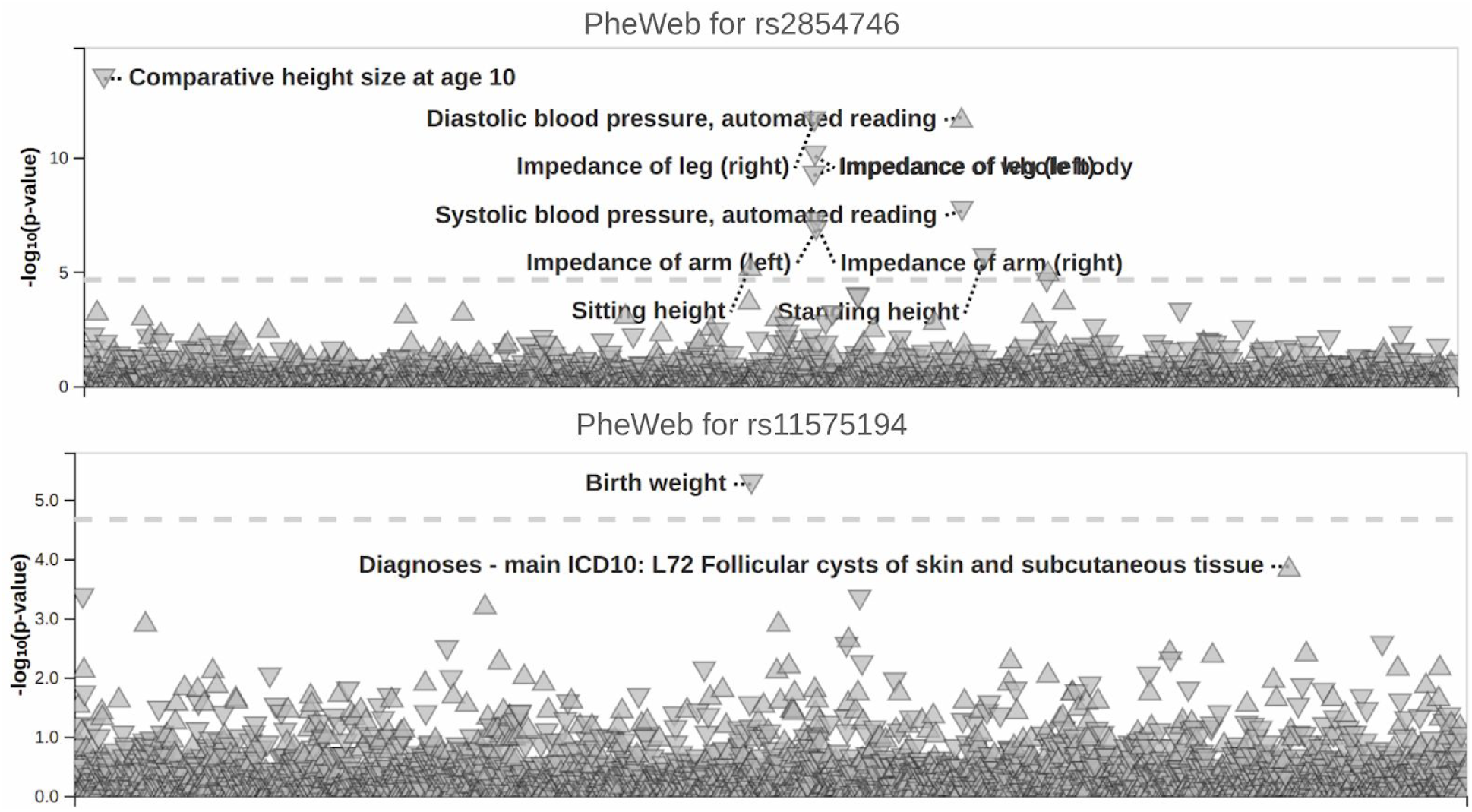
PheWeb lookup for primary signals. PheWeb for **A**. rs2854746. Comparative height size at age 10, standing and sitting height, as well as diastolic and systolic blood pressure and impedance of right and left arm and leg and of the whole body are all significantly associated with this SNP. Associations are far less significant than association with SHR. **B**. rs11575194. Birth weight is significantly associated.

**Supplemental Figure 4:**
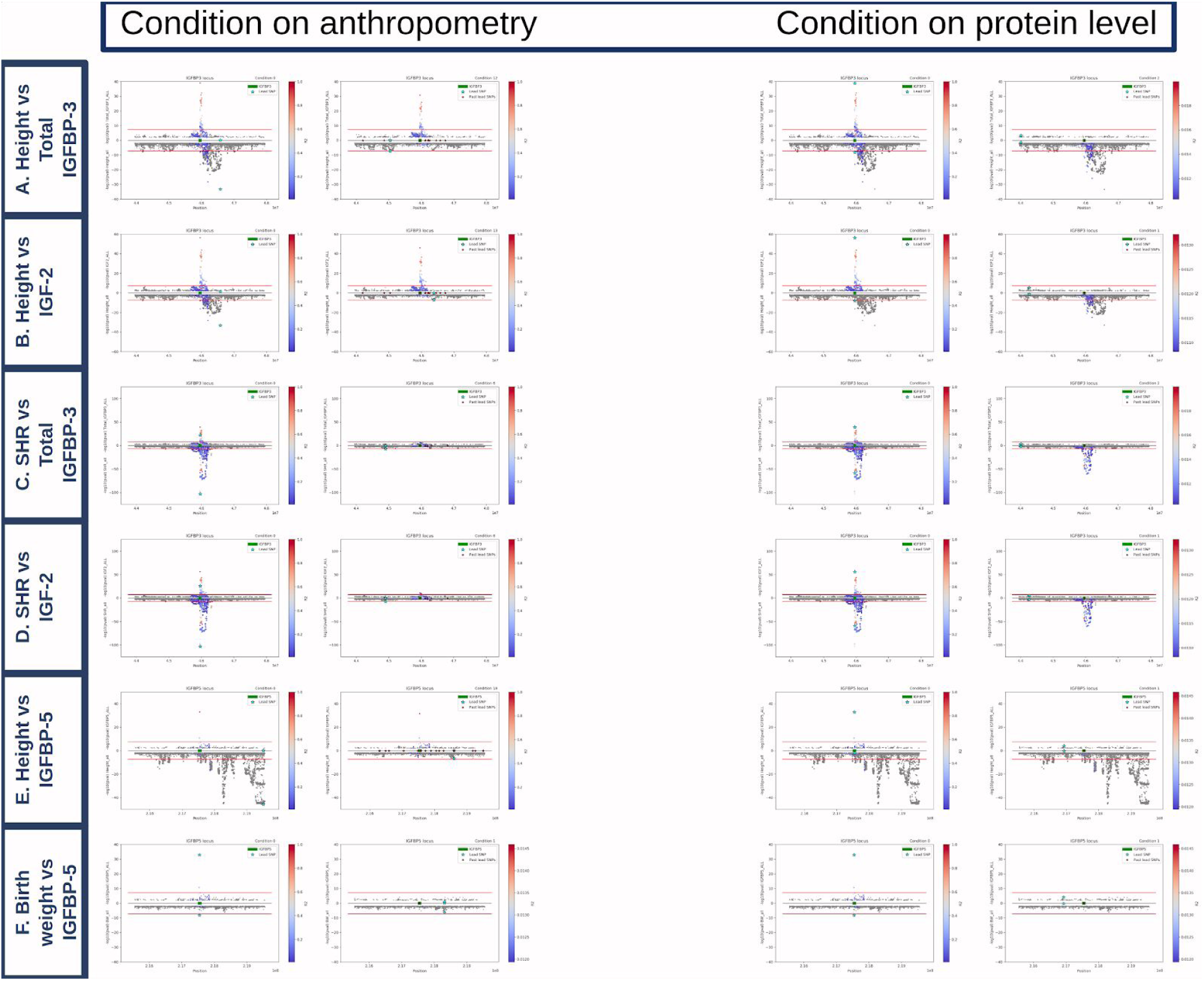
Conditional analyses. Extended from Figure 2. Left: conditioning on anthropometric associations in the region, Right: conditioning on serum protein level associations. For each pair: left: prior to conditioning; right: after all gws signal is conditioned away. When signal of protein level association remains after conditioning on anthropometry associations, or when signal of anthropometry association remains after conditioning on protein level associations; independence is concluded; when signal is attenuated, colocalization is concluded. Signal remains for height vs total IGFBP-3, height vs IGF-II, and height vs IGFBP-5 (both sides). Signal is attenuated for SHR vs total IGFBP-3, SHR vs IGF-II, and birth weight vs IGFBP-5 (both sides, especially pronounced when conditioning on anthropometry).

**Supplemental Figure 5:**
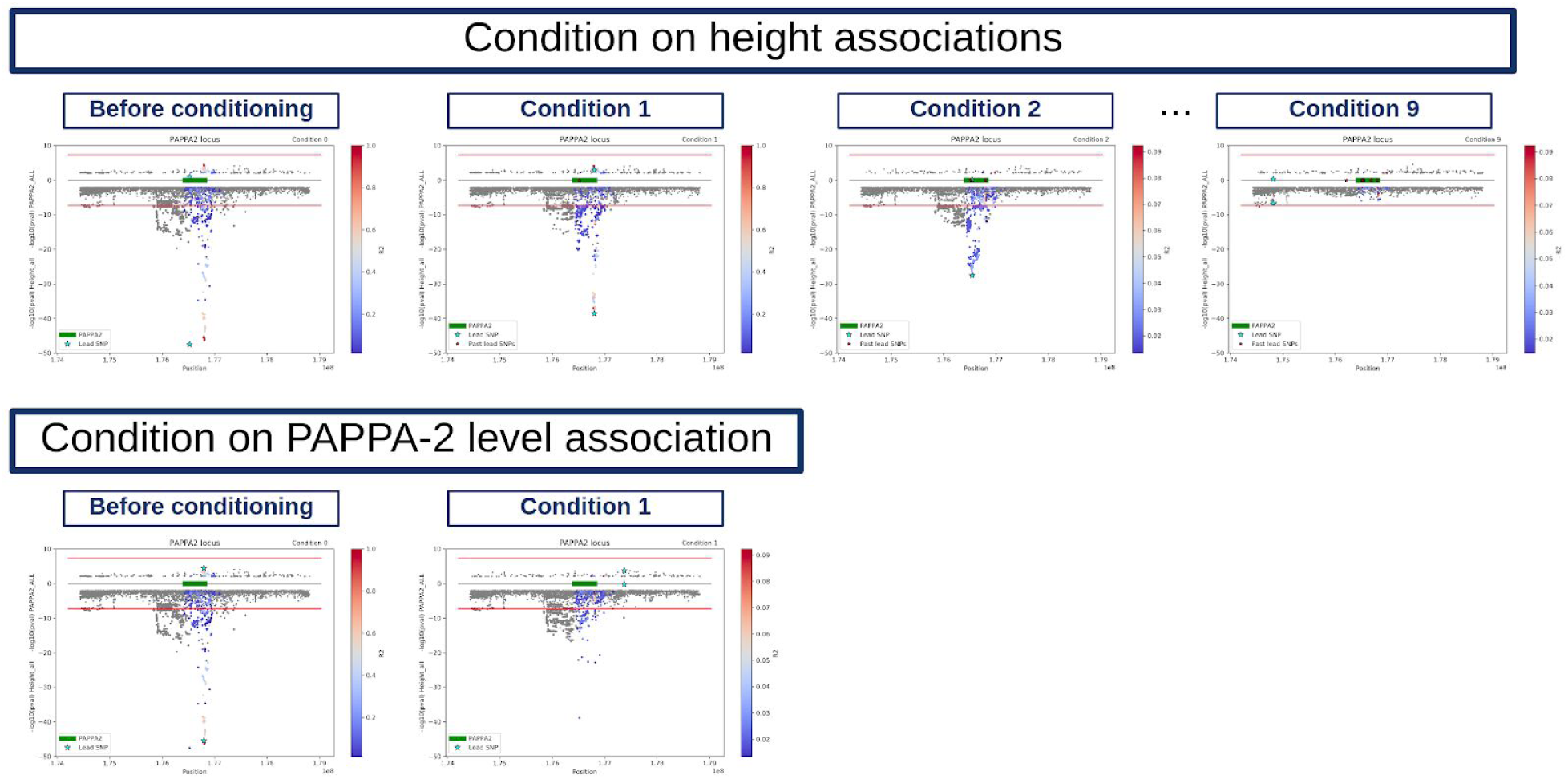
Conditional analysis, Height vs PAPP-A2. Top: conditioning on height associations in the region, bottom: conditioning on PAPP-A2 level associations. For each row: left: prior to conditioning; right: after all gws signal is conditioned away. In the top row, signal of association with PAPP-A2 levels is attenuated for the second condition, but not after the first condition. In the bottom row, regional height association is attenuated but not removed. We conclude that some height associations colocalize with nominal associations with PAPP-A2 levels, but some height associations are independent.

